# Structural insights into MBOAT2 catalysis, product retention, and ligand exchange

**DOI:** 10.64898/2026.09.05.749645

**Authors:** Jie Li, Jonathan Coria, Wook Shin, Lianqi Chen, Fangyu Liu

## Abstract

MBOAT2 suppresses ferroptosis independently of GPX4 and FSP1 by transferring monounsaturated acyl chains from acyl-CoA donors to lysophospholipid acceptors, but the structural basis of its catalytic cycle remains unclear. Here, we report the first cryo-EM structures of human MBOAT2, capturing endogenous and substrate-induced ligand-bound states. Unexpectedly, as-purified MBOAT2 contains a co-purified phospholipid-like density consistent with a retained product, together with a second density at a putative acyl-donor entry site. Oleoyl-CoA addition reduces the ordered product-like density and reveals donor density, whereas LPE addition increases local heterogeneity near the archway. The inactive H373A mutant contains endogenous donor- and acceptor-like densities along the two access pathways, consistent with substrate preloading. Together, these structures define the catalytic architecture of MBOAT2, support a product-retained, donor-primed working model, and provide templates for structure-guided ligand discovery.

## Introduction

Ferroptosis is a regulated form of cell death driven by iron-dependent phospholipid peroxidation. Its onset is governed by the balance between the generation of oxidizable membrane lipids and systems that limit lipid damage. GPX4 uses glutathione to reduce phospholipid hydroperoxides, whereas parallel pathways mediated by FSP1, DHODH, and GCH1 maintain radical-trapping metabolites that suppress lipid-peroxidation chain reactions^1–4^. Membrane composition provides an additional layer of control. Polyunsaturated fatty acid (PUFA)-containing phospholipids are particularly susceptible to peroxidation^5^. ACSL4 activates PUFAs, whereas LPCAT3/MBOAT5 incorporates them into membrane phospholipids, thereby promoting ferroptosis^5–7^. Conversely, enrichment of monounsaturated fatty acids (MUFAs) in phospholipids reduces membrane susceptibility to oxidation and protects cells from ferroptotic death^8^. Thus, ferroptosis can be suppressed not only by detoxifying lipid peroxides, but also by remodeling the membrane lipids in which peroxidation occurs.

MBOAT1 and MBOAT2 have recently emerged as key components of this protective lipid-remodeling pathway. These integral membrane acyltransferases suppress ferroptosis independently of GPX4 and FSP1 by incorporating monounsaturated acyl chains into phospholipids^9^. MBOAT2 is transcriptionally induced by androgen receptor signaling in prostate cancer and can preserve cell viability even in the absence of GPX4 and FSP1^9^. Beyond its role in ferroptosis resistance, elevated MBOAT2 expression has been reported in pancreatic cancer, where it is associated with aggressive disease and poor clinical outcome^10^. Together, these observations position MBOAT2 as a membrane-centered ferroptosis defense with broader relevance to cancer.

MBOAT2 is a member of <u>m</u>embrane-<u>b</u>ound <u>O</u>-<u>a</u>cyltransferases (MBOATs). MBOATs transfer acyl groups from acyl-coenzyme A (acyl-CoA) donors to lipid or protein acceptors and play critical roles in signal transduction and lipid metabolism. our phospholipid-remodeling MBOAT enzymes—MBOAT1, MBOAT2, MBOAT5 (LPCAT3), and MBOAT7 (LPIAT1)—catalyze the reacylation step of the Lands cycle by transferring fatty acyl groups from acyl-CoA donors to lysophospholipid acceptors^11–13^. Previous structural studies of LPCAT3/MBOAT5 and MBOAT7 have provided important insights into substrate recognition and selectivity^14,15^. Nevertheless, these enzymes differ in their phospholipid headgroup and acyl-chain preferences^15^. How MBOAT2 recognizes monounsaturated acyl-CoA donors and lysophospholipid acceptors, positions them for acyl transfer, and handles the resulting products remains unknown.

Here, we combine biochemical analysis and cryo-EM to define the catalytic architecture of human MBOAT2. Structures captured across endogenous and substrate-induced ligand states, together with structure-guided mutagenesis, reveal a preorganized intramembrane chamber and distinct access pathways for acyl-CoA and lysophospholipid substrates. These findings support a product-retained, donor-primed working model in which substrate-dependent ligand exchange resets MBOAT2 for subsequent catalysis.

## Results

### In vitro activity and overall structure of purified human MBOAT2

Human MBOAT2 was overexpressed and purified in detergent by affinity and size-exclusion chromatography. The purified protein eluted as two major populations, corresponding to monomeric and dimeric MBOAT2 (Extended Data Fig. 1a, b). To determine whether the purified enzyme retained catalytic activity, we compared purified wild-type (WT) MBOAT2 with the catalytically inactive H373A mutant. We established a continuous NADH-coupled fluorescence assay that measures acyltransferase activity through CoA production. WT MBOAT2 exhibited a Vmax of 773 nmol/mg/min and a Michaelis Menten constant (Km) of 22 μM for oleoyl-CoA (18:1-CoA) in the presence of 200 μM 18:1 lysophosphatidylethanolamine (LPE) (**Fig. 1a**), whereas MBOAT2 H373A exhibited minimal activity (**Fig. 1a**). The specific activity of purified human MBOAT2 was approximately significantly higher than that previously reported for mouse MBOAT2 in microsomal fractions and was comparable to related lipid-remodeling MBOAT enzymes, including MBOAT5 and MBOAT7^11,12,15,16^. Thus, purified MBOAT2 is a robust acyltransferase, consistent with its proposed ability to remodel membrane phospholipids in androgen-responsive ferroptosis defense^9^. We further profiled MBOAT2 activity across additional acyl-CoA donors. MBOAT2 strongly discriminated against the polyunsaturated donor arachidonoyl-CoA (20:4), exhibiting approximately 40-fold lower activity than with oleoyl-CoA (18:1) (**Fig. 1b**). In contrast, MBOAT2 processed palmitoleoyl-CoA (16:1) with activity comparable to that of oleoyl-CoA, consistent with previous reports^11^.

**Fig. 1:**
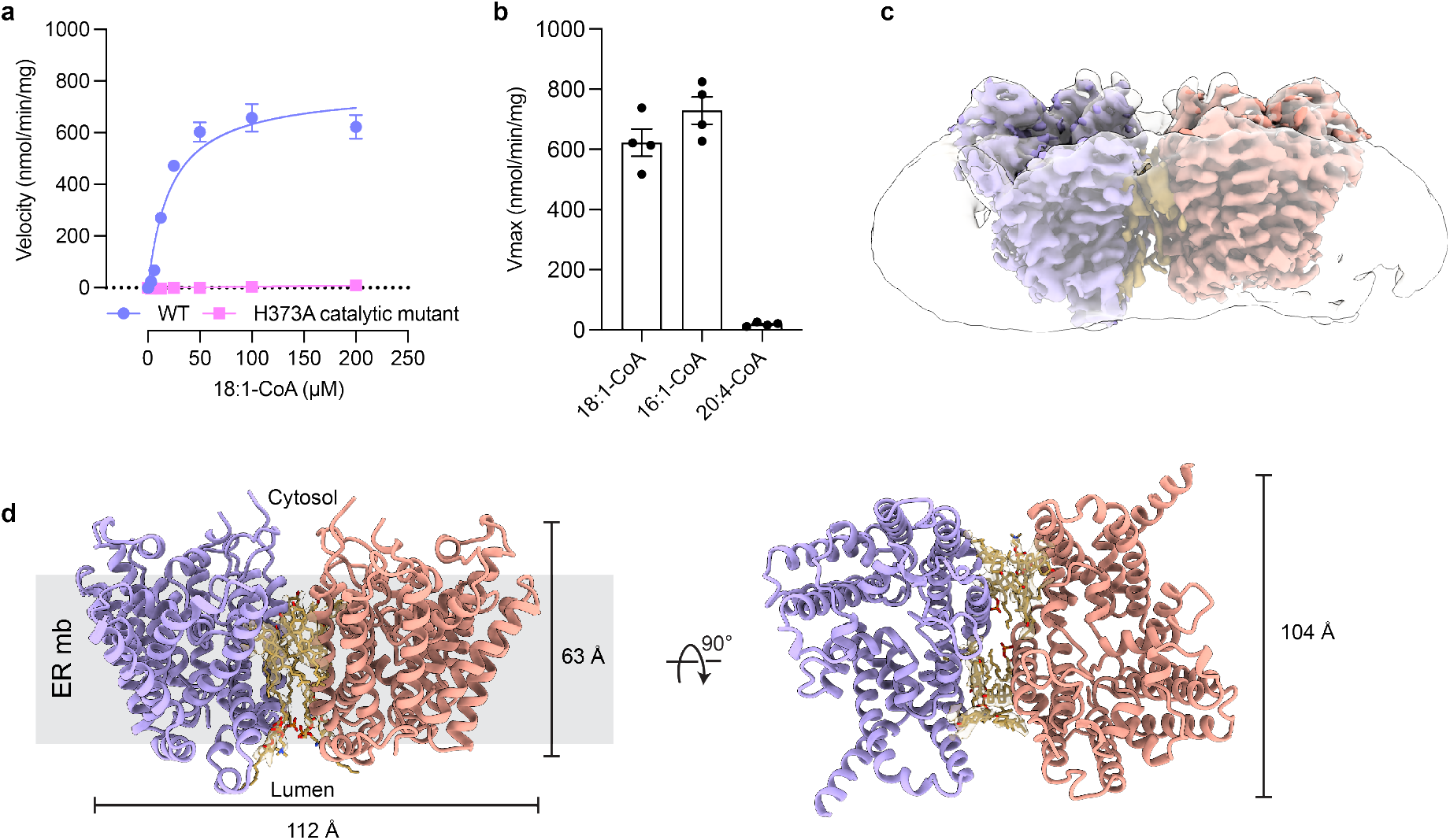
Biochemical characterization and overall structure of human MBOAT2. **a**, Enzymatic activity of wild-type human MBOAT2 and the catalytic mutant H373A. **b,** Acyl-donor selectivity of human MBOAT2 toward oleoyl-CoA (18:1-CoA), palmitoleoyl-CoA (16:1-CoA), and arachidonoyl-CoA (20:4-CoA). **c,** Cryo-EM density of the human MBOAT2 dimer embedded in detergent micelle. Protomer A and protomer B densities are colored lavender and coral, respectively. Densities for interfacial lipids are colored gold; the detergent micelle is shown in gray. **d,** Overall structure of the MBOAT2 dimer shown in cartoon representation, viewed from within the plane of the ER membrane and from the cytosolic side. Data are mean ± s.e.m. derived from four independent repeats.

We determined the structure of dimeric human MBOAT2 by single-particle cryo-electron microscopy (cryo-EM) at an overall resolution of 3.1 Å (Extended Data Table 1, Extended Data Fig. 2; Methods). The overall structure spans 112 Å in width and 63 Å in height (**Fig. 1c**). We fitted an AlphaFold model of MBOAT2 into each protomer and adjusted the model as needed (**Fig. 1d**). The dimer interface is predominantly lipid-mediated: the direct protein-protein interface is limited to 239 Å^2^ and is mediated by residues from TM6 of each protomer, whereas twelve modeled lipids expand the composite interface to 3,827 Å^2^ (Extended Data Fig. 3a). A lipid-mediated dimer was also reported for MBOAT5 (LPCAT3), which preferentially transfers arachidonoyl chains to LPLs^14^. Intriguingly, MBOAT2 and MBOAT5 form dimers through opposite faces of the MBOAT fold (Extended Data Fig. 3b). Why these enzymes adopt such different assemblies, and whether this difference influences their function in native membranes, remain to be determined.

### A co-purified PC/PE-like density occupies the MBOAT2 reaction chamber

Each MBOAT2 protomer comprises 11 membrane-spanning helices (TM1–TM11) connected by shorter α-helical segments and loops (Fig. 2a, Extended Data Fig. 4). It adopts a structural “MBOAT fold” similar to previously reported lipid- and protein-modifying MBOATs^17–24^.

**Fig. 2:**
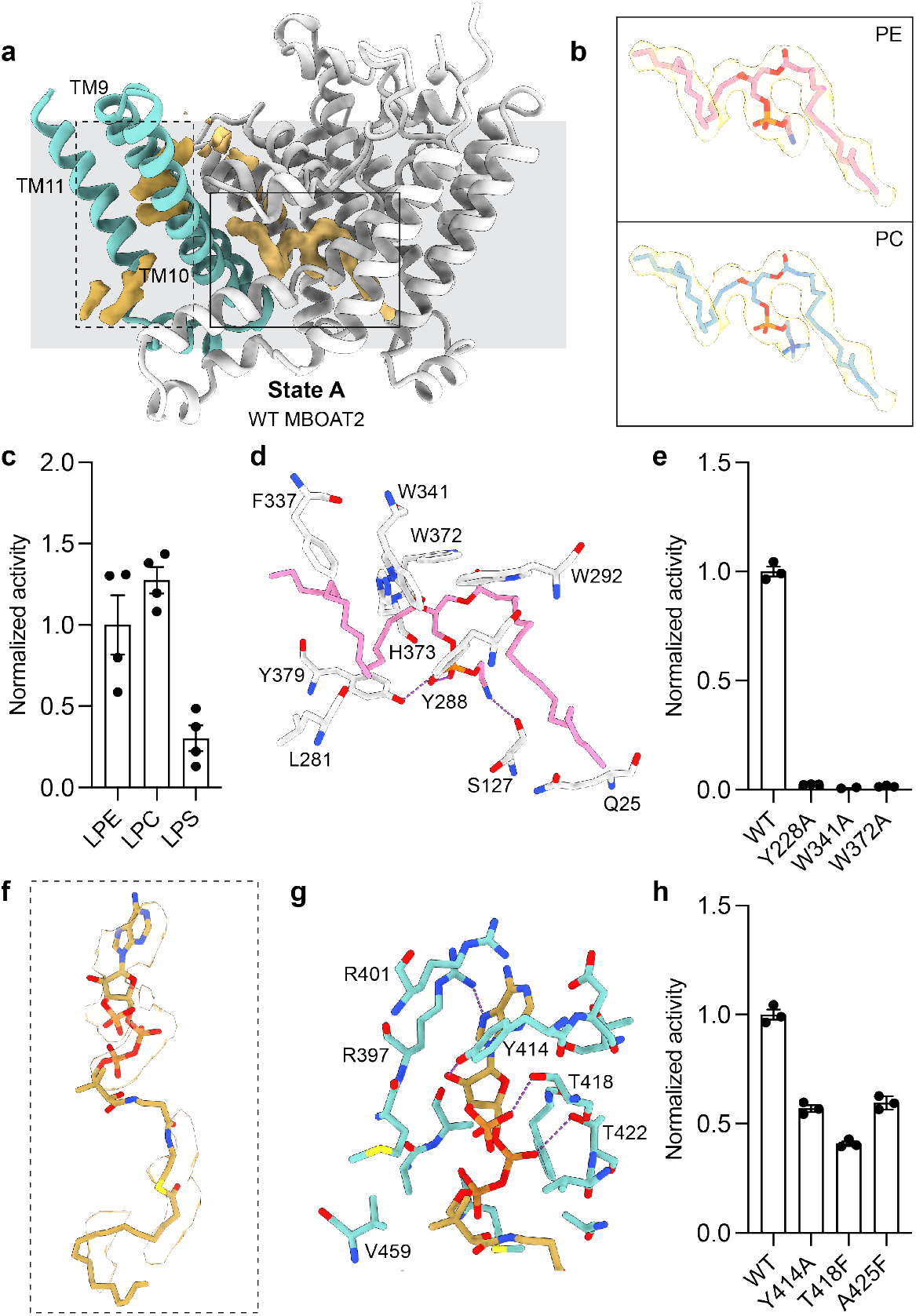
Co-purified ligand densities occupy the MBOAT2 reaction chamber and putative donor-entry site. **a**, Structure of monomeric MBOAT2 shown in ribbon representation. The protein is colored light gray, with the lateral-gate/archway region colored teal. Non-protein densities within the catalytic chamber and acyl-CoA gateway are shown in gold. **b,** Fitting of PE and PC into the observed catalytic-chamber density. **c,** Enzymatic activity of MBOAT2 toward LPE, LPC, and LPS acceptor substrates. **d,** Molecular interactions between the modeled phospholipid product and residues within 3.5 Å. **e,** Enzymatic activities of MBOAT2 mutants in the product-binding chamber. **f,** Zoomed-in view of the acyl-CoA gateway density, into which oleoyl-CoA was modeled. Oleoyl-CoA is shown in beige/gold. **g,** Molecular interactions between oleoyl-CoA and surrounding residues within 3.5 Å. **h,** Enzymatic activities of MBOAT2 mutants in the acyl-CoA gateway. Cryo-EM density surfaces were displayed at a contour level corresponding to 6 standard deviations above the mean map density, after local zoning around the modeled ligand. c, e, h, Data are mean ± s.e.m. derived from three independent repeats.

Inspection of the cryo-EM map revealed clear, non-protein densities within the reaction chamber of each MBOAT2 protomer (**Fig. 2a**). The density occupied the LPL-binding region and extended into the hydrophobic channel that accommodates the acyl chain of oleoyl-CoA, consistent with a phospholipid product-like assignment. Because no exogenous lipid was added during purification or grid preparation, this density likely represents a tightly associated endogenous lipid retained through detergent solubilization and purification.

The headgroup density is most compatible with PE or PC but did not permit an unambiguous chemical assignment (**Fig. 2b**, Extended Data Fig. 5). Purified MBOAT2 used both LPE and LPC as acceptors, whereas activity with LPS was much lower (**Fig. 2c**)^11,12,15^. Guided by the density, biochemical profile, and reported role of MBOAT2 in PE remodeling during ferroptosis^9^, we used dioleoyl phosphatidylethanolamine (DOPE) as a working model for subsequent structural interpretation (**Fig. 2d**). In this model, the phosphate headgroup forms hydrogen bonds with Y288 and Y379, whereas the positively charged ethanolamine moiety is positioned near the backbone carbonyl of S127. The acyl chains occupy a hydrophobic pocket lined by W292, F337, W341 and W372. Mutations of these pocket-lining residues markedly reduced MBOAT2 activity, showing that these residues are required for activity (**Fig. 2e**).

In addition to the well-defined phospholipid-like density, the map contained non-protein density along a putative acyl-donor entry pathway formed by TM9, 10 and 11 (**Fig. 2a, f**). This region forms an archway-like opening analogous to the putative palmitoyl-CoA recruitment site proposed for Hedgehog acyltransferase (HHAT), an MBOAT enzyme that acylates Hedgehog proteins^23,25^. The best-resolved portion of the density accommodated the CoA moiety of an acyl-CoA donor, suggesting a potential donor-docking site preceding entry into the reaction chamber. Mutations of residues lining this opening reduced, but did not abolish, oleoyl-CoA-dependent activity, consistent with a contribution to acyl-donor recruitment (**Fig. 2g, h**). Because the as-purified structure contains endogenous ligands and therefore does not represent a true apo state, we refer to it hereafter as **State A**.

### Oleoyl-CoA or LPE addition weakens the retained product-like density

We tested whether the co-purified phospholipid behaves as a retained product by examining how its density responds to added substrates. We incubated WT MBOAT2 with 200 μM oleoyl-CoA or 500 μM LPE for 30 min before grid freezing and determined structures at 3.4 and 3.5 Å, respectively, hereafter **State B** and **State C** (Extended Data Table 1, Extended Data Fig. 6, 7). Both ligand-added structures closely matched **State A**, with Cα r.m.s.d. values of 0.54 Å for State B and 0.60 Å for **State C**.

The State B reconstruction likely represents a mixture of oleoyl-CoA-bound and phospholipid-bound particles (**Fig. 3a, b**). Consistent with this interpretation, relative to **State A**, the ordered phospholipid density was reduced and accompanied by the appearance of oleoyl-CoA density (**Fig. 3b**). This reciprocal change is consistent with partial displacement of the co-purified phospholipid by the incoming acyl donor. Oleoyl-CoA is coordinated by salt bridges and hydrophobic interactions, and mutations within this pocket reduce catalytic activity (**Fig. 3c, d**).

**Fig. 3:**
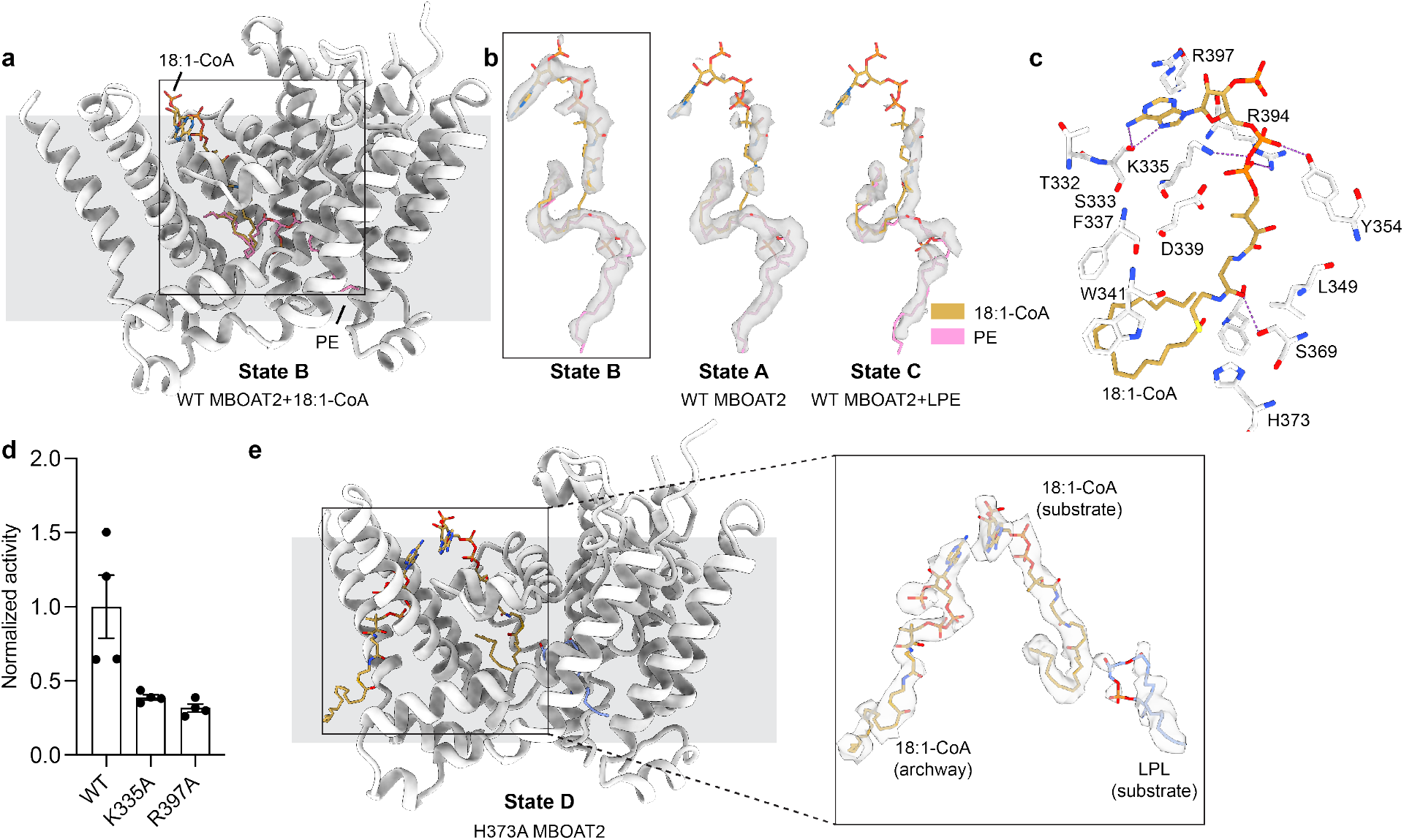
Substrate addition remodels ligand-associated density in MBOAT2. **a**, Structure of monomeric MBOAT2 determined after addition of 200 μM oleoyl-CoA, shown in ribbon representation. The protein is colored light gray. **b,** Comparison of cryo-EM densities within the catalytic chamber across different sample conditions. Oleoyl-CoA (18:1-CoA) is shown as yellow sticks and PE is shown as pink sticks, with corresponding ligand-associated densities shown as semi-transparent gray surfaces. PE-like and oleoyl-CoA models represent alternative members of an ensemble, not simultaneous ligands. **c,** Molecular interactions between oleoyl-CoA and its surrounding residues in the catalytic chamber. **d,** Enzymatic activities of MBOAT2 mutants targeting the oleoyl-CoA-binding site within the catalytic chamber. **e,** Structure of the catalytically inactive H373A MBOAT2 mutant shown in ribbon representation. Co-purified ligand-like densities are highlighted in the inset: oleoyl-CoA is shown as gold sticks and LPE is shown as light blue sticks. Ligand-associated cryo-EM densities were locally zoned around the modeled ligands and displayed at contour levels equivalent to 6 standard deviations above the mean density of the unzoned map.

To examine how acceptor binding affects the retained phospholipid, we determined **State C** after incubating WT MBOAT2 with 500 μM LPE. Rather than revealing a well-resolved LPE molecule, the map showed markedly weakened ordered phospholipid density (**Fig. 3b**). The residual density was weak and discontinuous, precluding confident modeling of a single ligand and suggesting that this condition populates a heterogeneous ligand-exchange ensemble. The archway-forming region was also less well resolved than in the other states, consistent with increased local heterogeneity (Extended Data Fig. 8). These observations raise the possibility that LPE engagement promotes local archway dynamics that facilitate oleoyl-CoA entry into the reaction chamber.

### Catalytic mutant captures substrate-like densities

If State A represents a product-retained conformation, we asked which ligands would be captured when catalysis was disrupted. We therefore determined the structure of the catalytically inactive H373A mutant without adding exogenous ligands, hereafter **State D** (**Fig. 3b**, Extended Data Table. 1, Extended Data Fig. 9). H373A retained the overall MBOAT2 fold, with a Cα r.m.s.d. of 0.77 Å relative to **State A**, indicating that the mutation did not globally disrupt the protein. The reaction chamber contained strong acyl-CoA-like density that closely resembled the donor density in **State B** and was modeled as oleoyl-CoA. Unlike **State A**, **State D** lacked an ordered phospholipid headgroup. Strong hydrophobic density nevertheless occupied the region expected to accommodate the sn-1 acyl chain of a lysophospholipid acceptor. This pattern suggests that acceptor-tail engagement can occur without a well-ordered headgroup.

### A structure-informed working model for MBOAT2 catalysis

Our biochemical and structural data support a structure-informed working model in which ligand exchange occurs within a largely preorganized reaction chamber (**Fig. 4**). In the state captured after purification, a phospholipid product occupies the chamber while an acyl-CoA donor is positioned at the cytosolic archway. Product departure may allow this pre-docked donor to advance into the acyl-donor pocket. Alternatively, LPE entry from the ER lumen is associated with increased local heterogeneity around the lateral gate, potentially promoting product exchange and donor entry. Because mutations of archway-lining residues reduced but did not abolish catalysis, an additional route for direct acyl-CoA access from the cytosol remains possible. These pathways would converge on a complex in which oleoyl-CoA and LPE are positioned near the catalytic histidine. Following acyl transfer, CoA is proposed to dissociate while the newly formed phospholipid remains associated with the chamber, regenerating the product-retained state.

**Fig. 4:**
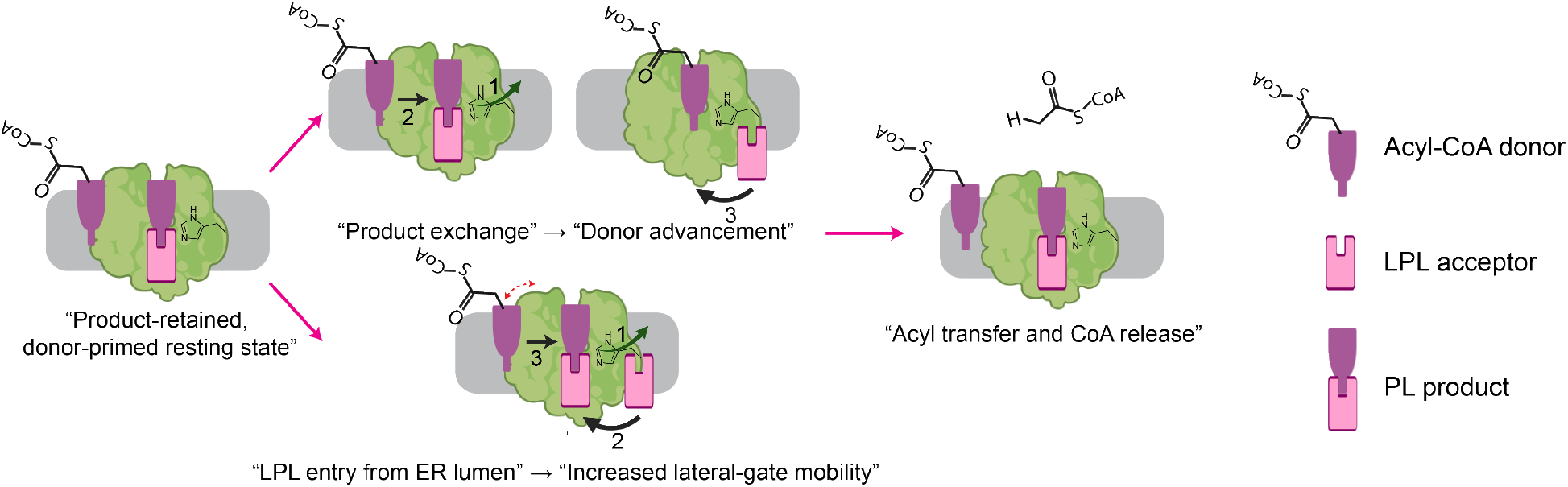
Proposed working model for MBOAT2 catalysis. The state captured in the as-purified sample contains a modeled phospholipid product within the reaction chamber and a donor-like species at the cytosolic archway. In the upper pathway, product dissociation (1) permits the pre-docked donor to advance into the acyl-donor pocket (2), followed by LPL acceptor entry from the ER lumen. In the lower pathway, LPL acceptor engages the acceptor pathway before product release and is associated with increased local heterogeneity around the lateral gate and archway, potentially facilitating product exchange and donor entry (3). Both pathways would converge on a catalytically competent complex in which oleoyl-CoA and LPL are positioned near His373. Acyl transfer is proposed to release CoA while the newly formed phospholipid remains associated with the reaction chamber, regenerating the product-retained state. Direct acyl-CoA entry from the cytosol is possible but is not illustrated. The red dashed arrow denotes increased local mobility. LPL: lysophospholipid. PL: phospholipid.

## Discussion

Our structural and biochemical analyses lead to three principal conclusions. First, the as-purified MBOAT2 structure captures a stable endogenous phospholipid-like state. In an LPC-supplemented LPCAT3/MBOAT5 structure, an unmodeled density was proposed to represent partially co-purified PC, but its identity and relationship to catalysis remained unresolved^14^. In MBOAT2, the ligand forms continuous density across the LPL/product-binding region and extends into the acyl-chain channel. Its ordered density decreased following oleoyl-CoA addition, its headgroup density was absent in the catalytically inactive H373A mutant, and its ordered density was weakened by LPE supplementation. Together, these observations support a working interpretation of the ligand as a retained phospholipid product and provide a systematic structural characterization of a product-like state in a lipid-remodeling MBOAT.

Second, the captured ligand states identify product exchange as a likely step in catalytic resetting and reveal how ligand occupancy reshapes the MBOAT2 reaction chamber. The protein scaffold remains largely unchanged across these states, whereas ligand density and local order change at the proposed access pathways. Catalytic cycling therefore appears to involve local ligand exchange and gate dynamics within a preorganized chamber. Occupancy by the donor or product substantially reduces the accessible chamber volume and generates smaller, state-specific pockets. Moreover, the persistence of endogenous donor- and product-like species through detergent solubilization and multistep purification is compatible with tight or slow-exchange binding, although the structures cannot distinguish thermodynamic affinity from kinetic stability. These conformations may support the development of state-selective MBOAT2 modulators and, if conserved, could reveal ligandable states across the broader MBOAT family.

Third, these first experimental structures of MBOAT2 provide a framework for understanding its substrate selectivity. Purified MBOAT2 efficiently utilized oleoyl-CoA and palmitoleoyl-CoA while strongly discriminating against arachidonoyl-CoA. The preference of MBOAT2 for monounsaturated donors is consistent with its role in enriching membranes with phospholipids that are less susceptible to peroxidation. The lipid-mediated MBOAT2 dimer represents an additional structural feature, although whether distinct MBOAT assemblies influence activity, localization, or lipid access in native membranes remains unknown.

Several aspects of the proposed working model remain unresolved. The structures do not establish the residence time or release pathway of the retained product. Likewise, increased archway mobility following LPE addition is inferred from weakened local density, and residual activity of archway mutants does not establish an alternative donor-entry route. Resolving these questions will require direct kinetic measurements and additional ligand-resolved intermediates. Together, our findings define the catalytic architecture of MBOAT2 and identify persistent ligand-associated conformations as mechanistically informative and potentially ligandable states.

## Materials and Methods

### Protein expression and purification

The cDNA encoding full-length human MBOAT2 (UniProt accession Q6ZWT7) was cloned into the pEG BacMam expression vector^26^ with a C-terminal green fluorescent protein (GFP) tag. A PreScission protease cleavage site was introduced between MBOAT2 and GFP to permit removal of GFP during purification. Recombinant proteins were expressed by baculovirus-mediated transduction of HEK293S GnTI− cells (ATCC) following an established protocol^26^.

To purify the protein, cells expressing MBOAT2 were resuspended in Buffer A containing 20 mM HEPES, pH 7.5, 200 mM NaCl, 20% glycerol, 5 mM MgCl2, 1% 2,2-didecylpropane-1,3-bis-β-D-maltopyranoside (LMNG), 5 µg/mL DNase and protease inhibitors (1 μg/ml leupeptin, 1 μg/ml pepstatin, 1 μg/ml aprotinin, 100 μg/ml trypsin inhibitor, 1 mM benzamidine). The suspension was incubated for 2 h at 4 °C with rotation. Insoluble material was removed by centrifugation at 60,000 × g for 40 min at 4 °C. The clarified supernatant was incubated for 1.5 h at 4 °C with 2 mL of GFP nanobody-conjugated Sepharose resin that had been pre-equilibrated with Buffer A.

The resin was washed with five column volumes of buffer containing 20 mM HEPES, pH 7.5, 200 mM NaCl, 2 mM dithiothreitol and 0.01% glyco-diosgenin (GDN). MBOAT2 was released by on-resin cleavage with GST-tagged PreScission protease for 2 h at 4 °C, and the eluate was passed over glutathione resin to remove the protease. The protein was concentrated and purified by size-exclusion chromatography on a Superose 6 Increase 10/300 GL column.

### In vitro acylation activity assay

MBOAT2 activity was measured with a fluorescence-based coupled-enzyme assay on a CLARIOstar Plus plate reader^27,28^. Reactions were carried out in an assay buffer with 20 mM HEPES, pH 7.5, 150 mM NaCl, 2 mM β-mercaptoethanol, 0.01% GDN. The reaction mixture contained 0.25 mM NAD+, 0.2 mM thiamine pyrophosphate and 2 mM α-ketoglutarate. α-Ketoglutarate dehydrogenase (αKDH) was purified from bovine heart obtained from a local meat supplier according to a previously described procedure^29^. An appropriate amount of αKDH was used to ensure that the MBOAT2 reaction is the rate-limiting step.

For acyl-CoA kinetics, 18:1 lyso-PE was maintained at 200 µM while acyl-CoA was varied from 0 to 200 μM. For lysophospholipid kinetics, oleoyl-CoA was maintained at 200 µM while lysophospholipids were varied from 0 to 200 µM. Data were fitted to the Michaelis–Menten equation using GraphPad Prism 10.

### Cryo-EM sample preparation and data collection

Quantifoil R1.2/1.3 400-mesh Au holey carbon grids (Quantifoil) were glow-discharged for 30 s. Protein samples were concentrated to 5 to 10 mg/mL, supplemented with 0.1 mM FF8, and applied to the glow-discharged grids. The grids were blotted for 3.5 s using a Vitrobot (FEI) and plunge-frozen in liquid ethane cooled by liquid nitrogen. For substrate-supplemented MBOAT2 samples, the protein was incubated on ice for 30 min with either 500 μM 18:1 lyso-PE or 200 μM oleoyl-CoA and then centrifuged for 5 min before grid preparation.

Cryo-EM data were collected using SerialEM on a Titan Krios microscope (FEI) operated at 300 kV. All datasets were collected using a Falcon 4i detector at a calibrated pixel size of 0.94 Å per pixel and the same nominal defocus range of −1.8 to −0.8 μm. Each movie was recorded over 4 s with a total electron dose of approximately 40 e⁻ Å⁻².

### Cryo-EM data processing

All cryo-EM data processing was performed using cryoSPARC v4.0^30^. Raw movie stacks were subjected to motion correction using MotionCor2^31^, followed by estimation of contrast transfer function (CTF) parameters using Patch CTF estimation.

For the initial WT MBOAT2 dataset (dataset 1), 3,135 raw movie stacks were processed, from which 650,278 particles were extracted and subjected to multiple rounds of 2D classification. Particles from well-defined 2D classes were selected for ab initio reconstruction, followed by heterogeneous refinement to separate structurally distinct and poorly resolved particle populations. The particle subset displaying the best-defined MBOAT2 dimer features was subjected to an additional round of ab initio reconstruction and heterogeneous refinement, yielding a reconstruction at 5.16 Å resolution. Subsequent refinement improved the overall resolution to 4.78 Å. This map was then used as an initial reference volume for processing the larger WT MBOAT2 dataset (dataset 2). A total of 7,852 raw movie stacks were motion-corrected and CTF-estimated as described above. A total of 5,673,955 particles were initially extracted and subjected to 2D classification. Particles corresponding to well-resolved MBOAT2 views were selected for 3D classification by ab initio reconstruction and heterogeneous refinement. The particle subset exhibiting well-defined structural features was subjected to non-uniform refinement, followed by local refinement, resulting in a final reconstruction at an overall resolution of 3.14 Å.

For the oleoyl-CoA- and 18:1 LPE-supplemented complexes, dark-subtracted images were first normalized using the gain reference at a calibrated pixel size of 0.94 Å per pixel. Motion correction and CTF estimation were performed in cryoSPARC v4.4.1. Particles were automatically picked using templates generated from the WT MBOAT2 reconstruction, and the particle picks were manually inspected before extraction. Particles were initially extracted in a 320-pixel box and Fourier-cropped to 64 pixels. Subsequent processing followed the same general workflow described for the WT MBOAT2 datasets, including non-uniform refinement, local refinement, and focused 3D classification using a mask that excluded the detergent micelle density. Particles from the best-resolved classes were re-extracted in a 320-pixel box and subjected to further refinement to obtain the final reconstructions.

For the H373A mutant dataset, 8,745 movies were subjected to motion correction using MotionCor2 and CTF estimation using Patch CTF, resulting in 8,453 micrographs for further processing. Template-based particle picking and manual inspection yielded 3,824,316 extracted particles. Following 2D classification, 2,162,666 particles were retained and subjected to ab initio reconstruction and two rounds of heterogeneous refinement. The resulting particle subset was re-extracted at bin 2 and subjected to non-uniform refinement. Particles were then re-extracted at bin 1 and further processed by ab initio reconstruction, heterogeneous refinement, and non-uniform refinement. Finally, 325,963 particles were selected for focused refinement, yielding a final reconstruction at 3.54 Å resolution. The resolutions of all final maps were estimated using the gold-standard Fourier shell correlation criterion at an FSC cutoff of 0.143^32^.

### Model building and refinement

An AlphaFold-predicted model was fitted into the WT MBOAT2 cryo-EM map and manually adjusted in Coot^33^. The refined WT MBOAT2 model was subsequently fitted into the maps of the oleoyl-CoA-supplied MBOAT2 complex, the 18:1 LPE-supplied MBOAT2 complex, and the H373A MBOAT2 mutant, followed by manual adjustment in Coot^33^. Protein residues and well-resolved lipids were modeled according to the corresponding cryo-EM densities. All models were refined in real space using PHENIX^34^ with secondary-structure and geometry restraints and validated using MolProbity^35^. Structural figures were prepared using UCSF ChimeraX. Model refinement and validation statistics are summarized in Extended Data Table 1.

## Acknowledgments

Cryo-EM data were collected at the UT Southwestern Medical Center Cryo-Electron Microscopy Facility. We thank Xiaochun Li, Xuewu Zhang and Boyuan Wang for helpful discussions, and Fenfang Yang and Xin Li for experimental advice. This work was supported by awards from the Cancer Prevention and Research Institute of Texas (CPRIT; RR250017) and the Damon Runyon Cancer Research Foundation (72-25) to F.L.

## Author contributions

J.L. and F.L. designed research; J.L., J.C., W.S. and L.C. performed research.

## Data availability

The 3D cryo-EM maps have been deposited in the Electron Microscopy Data Bank under the accession numbers EMD-XXXXX. Atomic coordinates for the atomic model have been deposited in the Protein Data Bank under the accession numbers XXXX.

