## Supplemental information for "Structural insights into MBOAT2 catalysis, product retention, and ligand exchange"

**This PDF file includes:**
Extended Data Fig. 1 to 9
Extended Data Table 1
Supplementary References

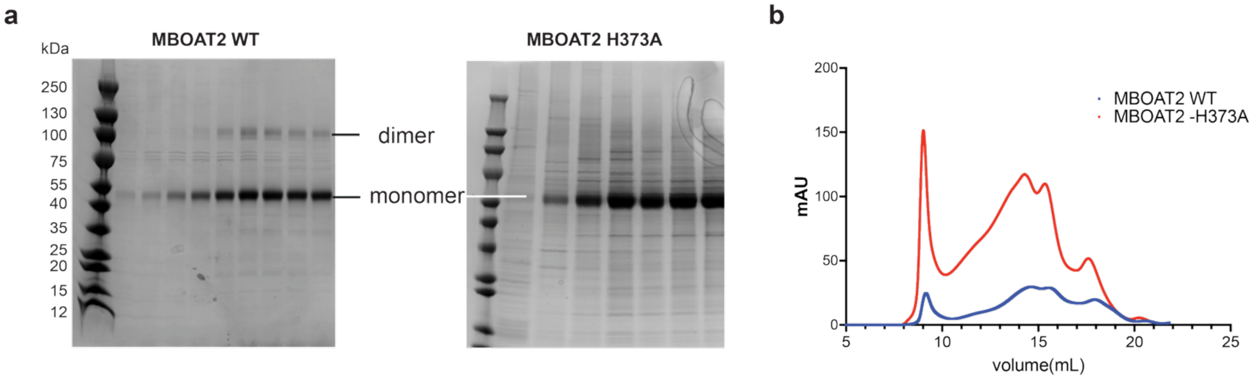


**Extended Data Fig. 1: Purification of human MBOAT2. a,** SDS-PAGE analysis of purified wild-type MBOAT2 and the catalytic mutant H373A. **b,** Size-exclusion chromatography profile of purified human MBOAT2. MBOAT2 eluted as two major peaks: an earlier peak at approximately 14 ml, corresponding to the dimeric species, and a later peak at approximately 15.2 ml, corresponding to the monomeric species.

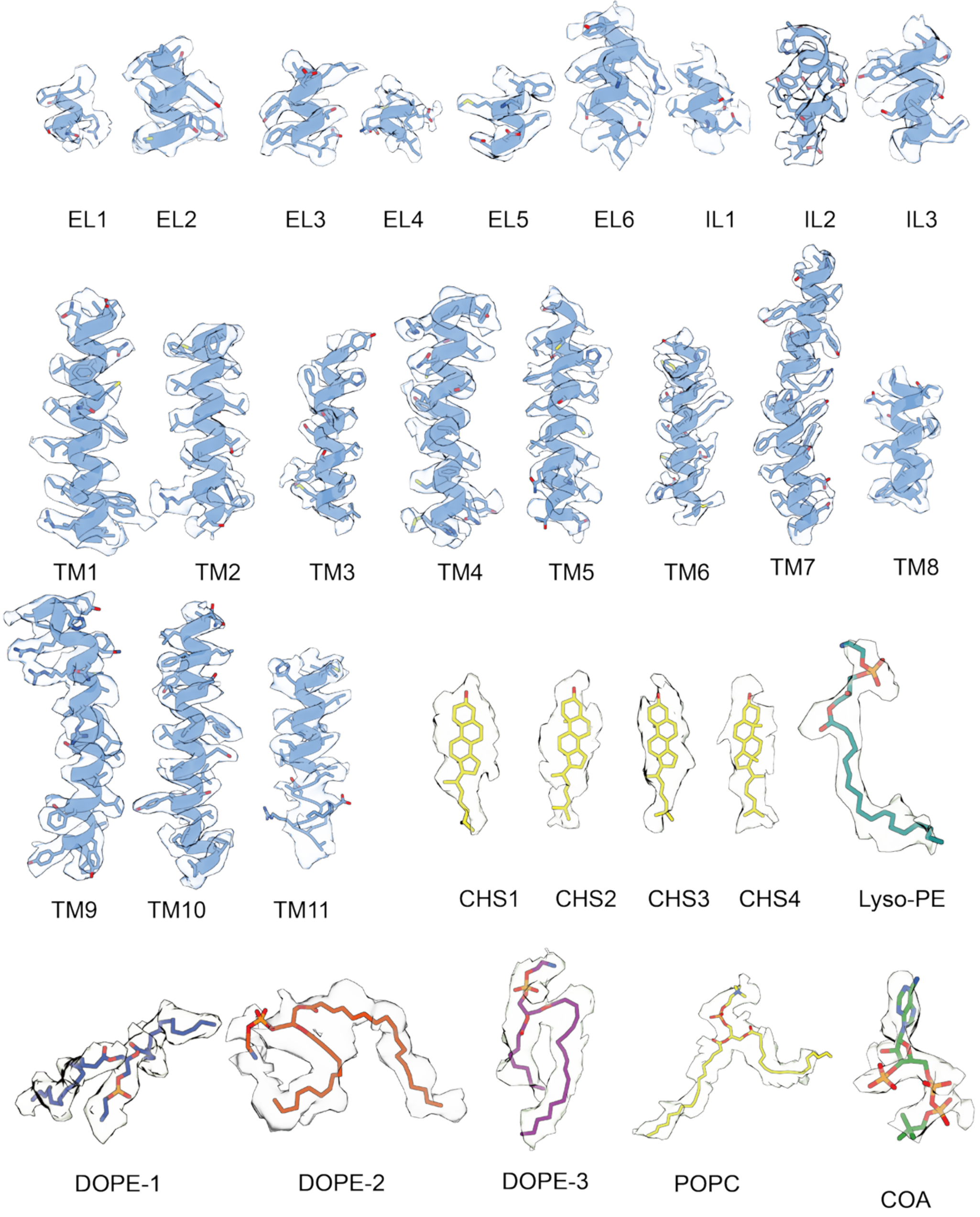


**Extended Data Fig. 2: EM density of different parts of human MBOAT2 in State A.**

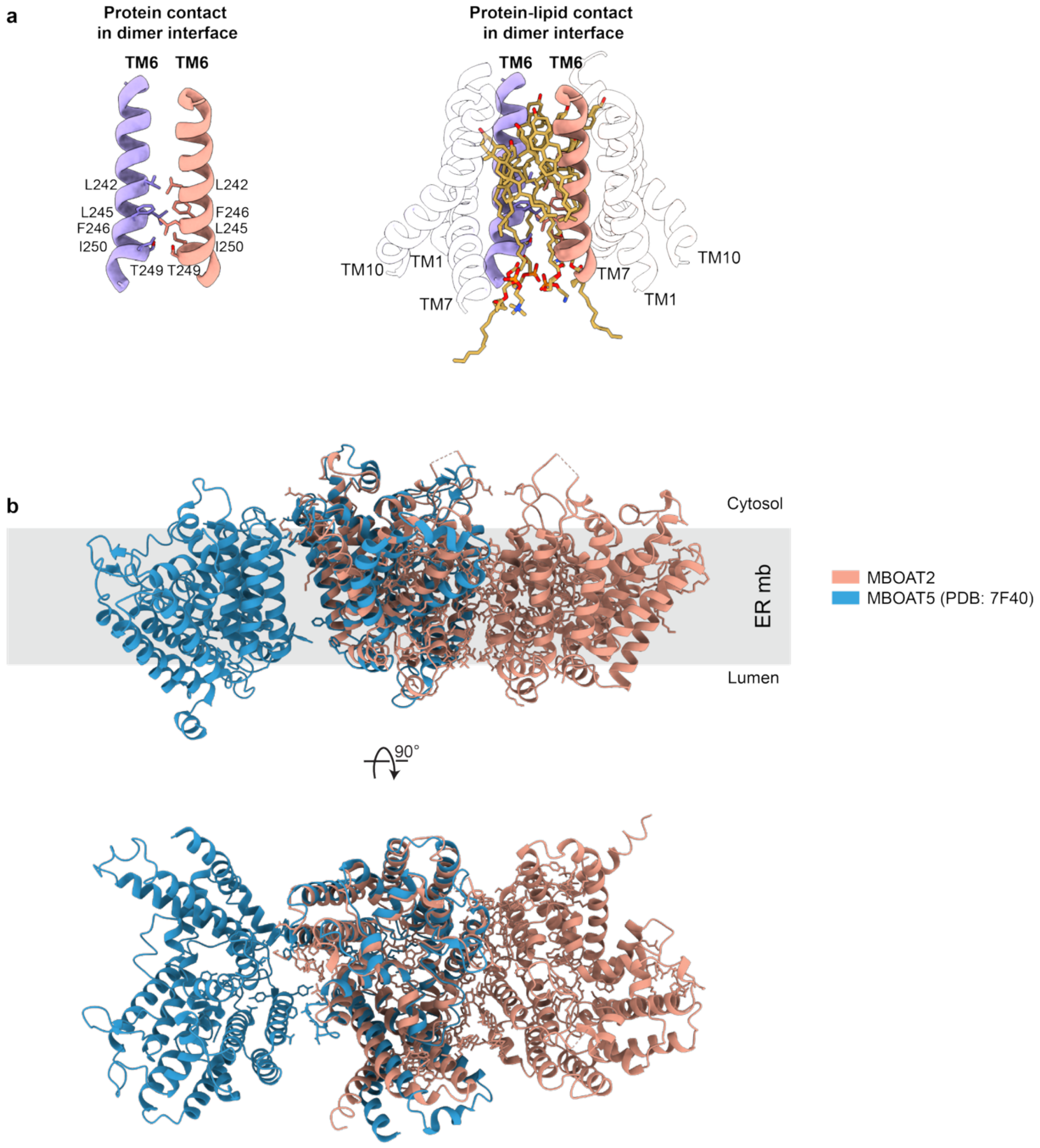


**Extended Data Fig. 3: Structural details of the human MBOAT2 dimer interface. a,** Close-up view of the direct protein-protein interface between MBOAT2 protomers. The interface is mediated primarily by TM6 from each protomer, with hydrophobic contacts contributed by L242, L245, F246, T249, and I250. **b,** Structural comparison of the MBOAT2 and MBOAT5 dimers[1].

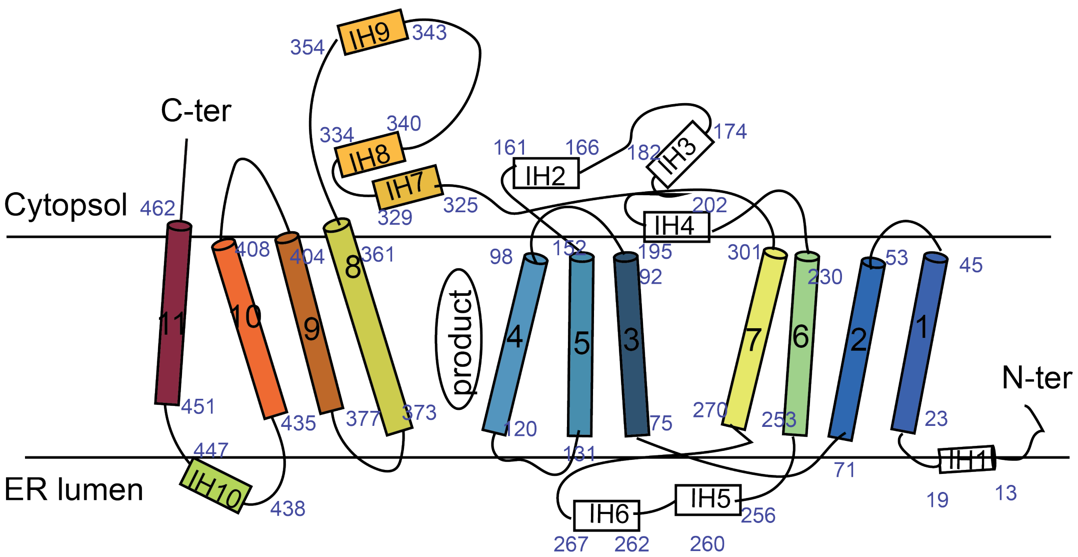


**Extended Data Fig. 4: The domain structure of human MBOAT2.**

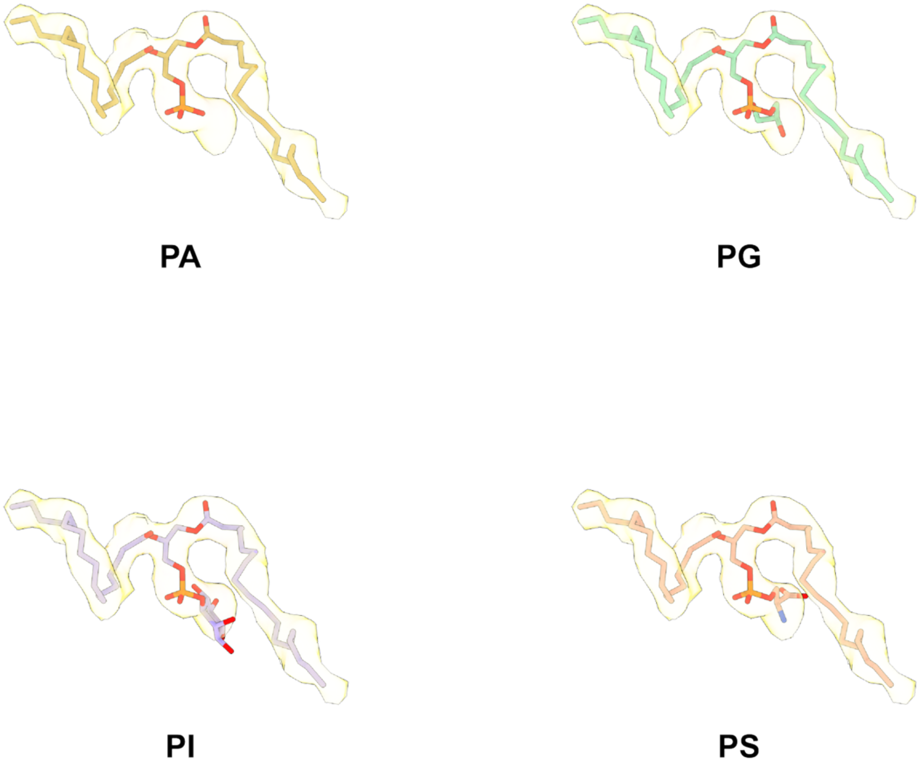


**Extended Data Fig. 5: Fitting of alternative phospholipids into the catalytic-chamber density.** Alternative phospholipid headgroups were modeled into the catalytic-chamber density.

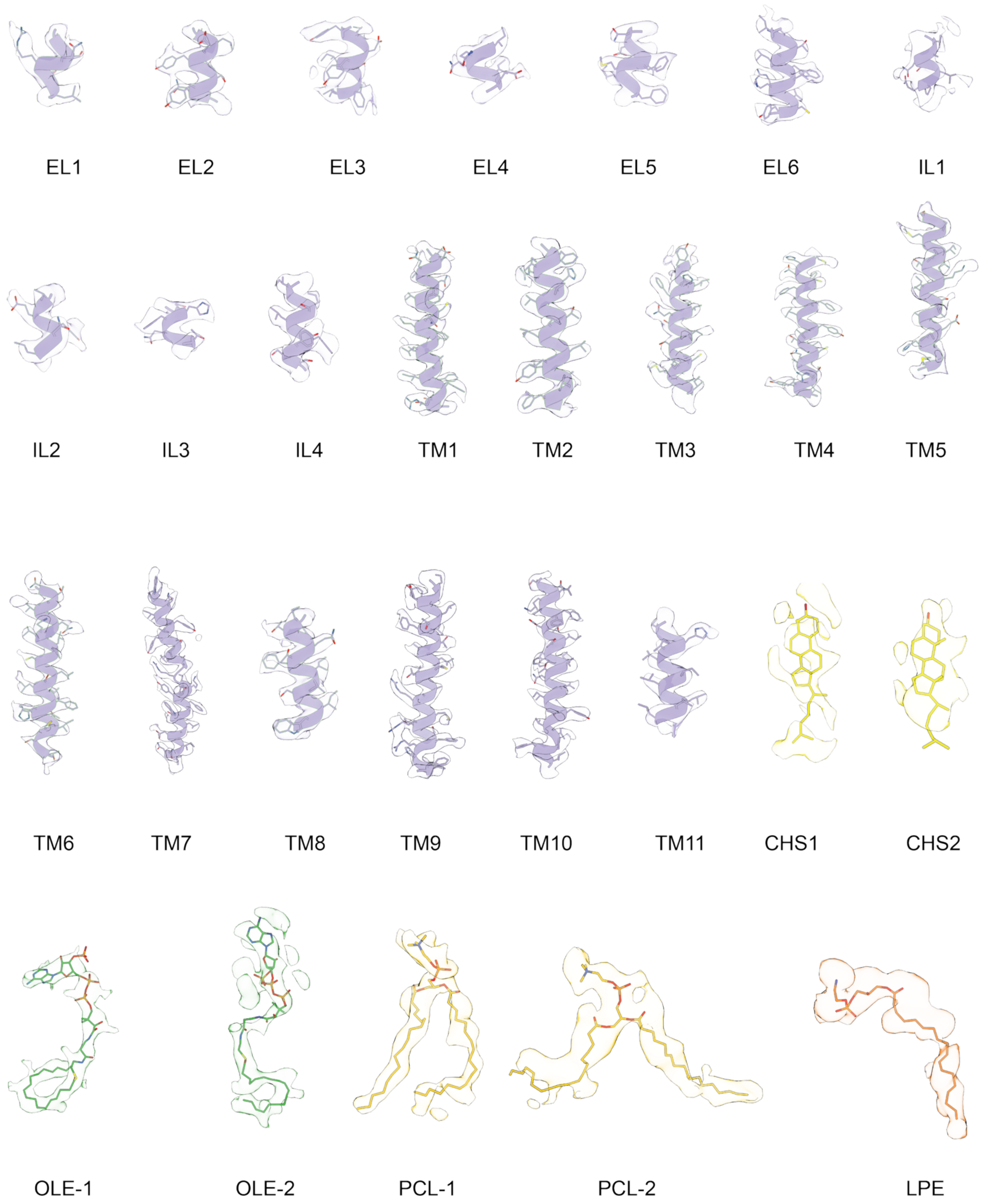


**Extended Data Fig. 6: EM density of different parts of human MBOAT2 in State B.**

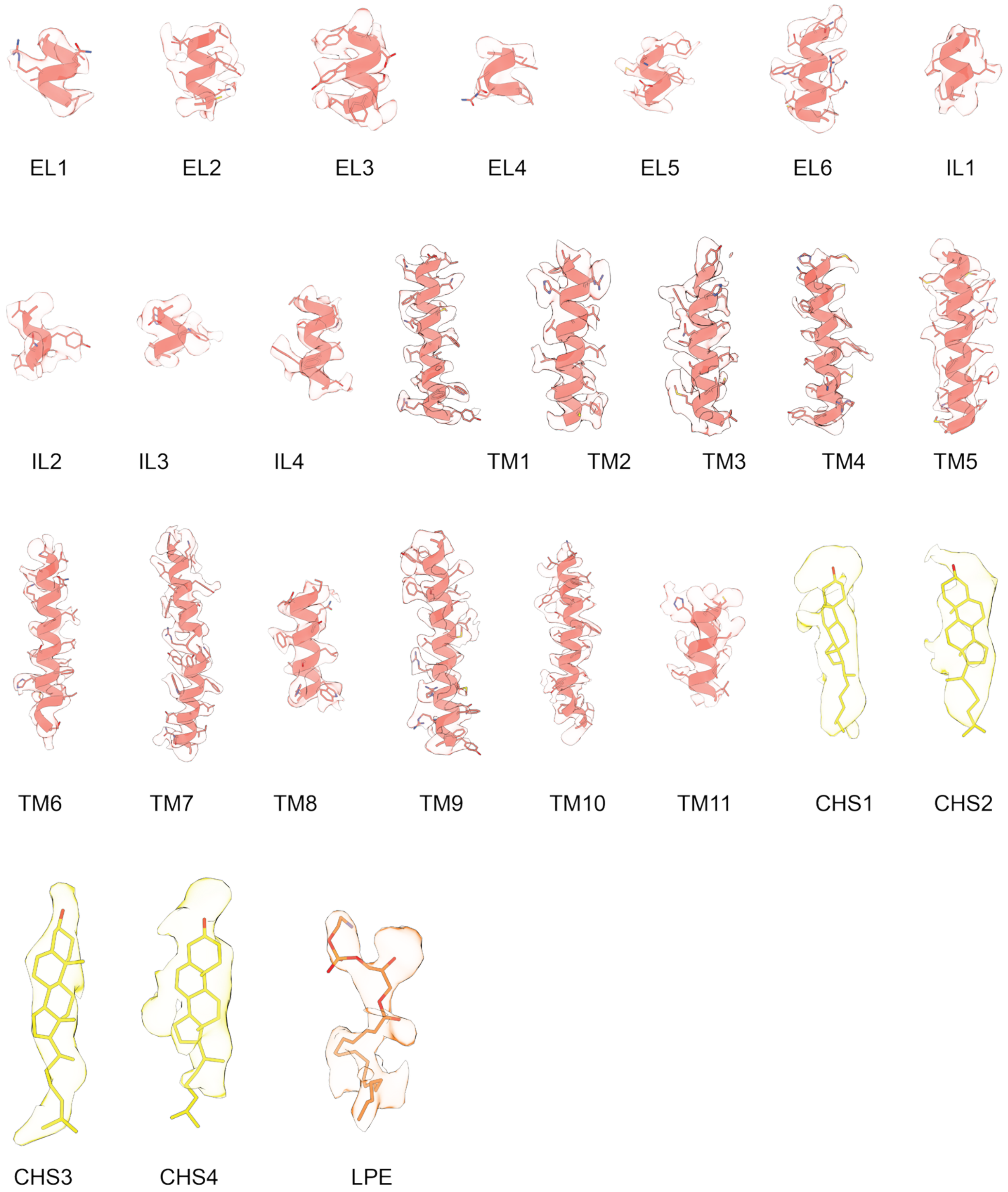


**Extended Data Fig. 7: EM density of different parts of human MBOAT2 in State C.**

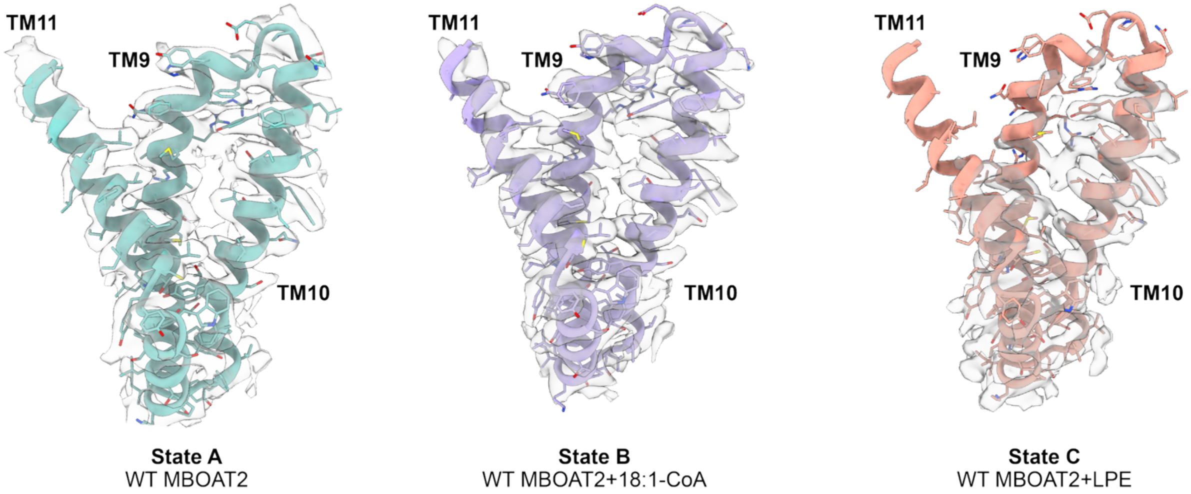


**Extended Data Fig. 8: Comparison of local mobility in the putative substrate-entry archway.** Cryo-EM densities surrounding the putative substrate-entry archway are compared across MBOAT2 states. Maps were locally zoned around the archway region and displayed at contour levels equivalent to 6 standard deviations above the mean density of the corresponding unzoned map. Reduced or fragmented density in this region indicates increased local mobility or conformational heterogeneity.

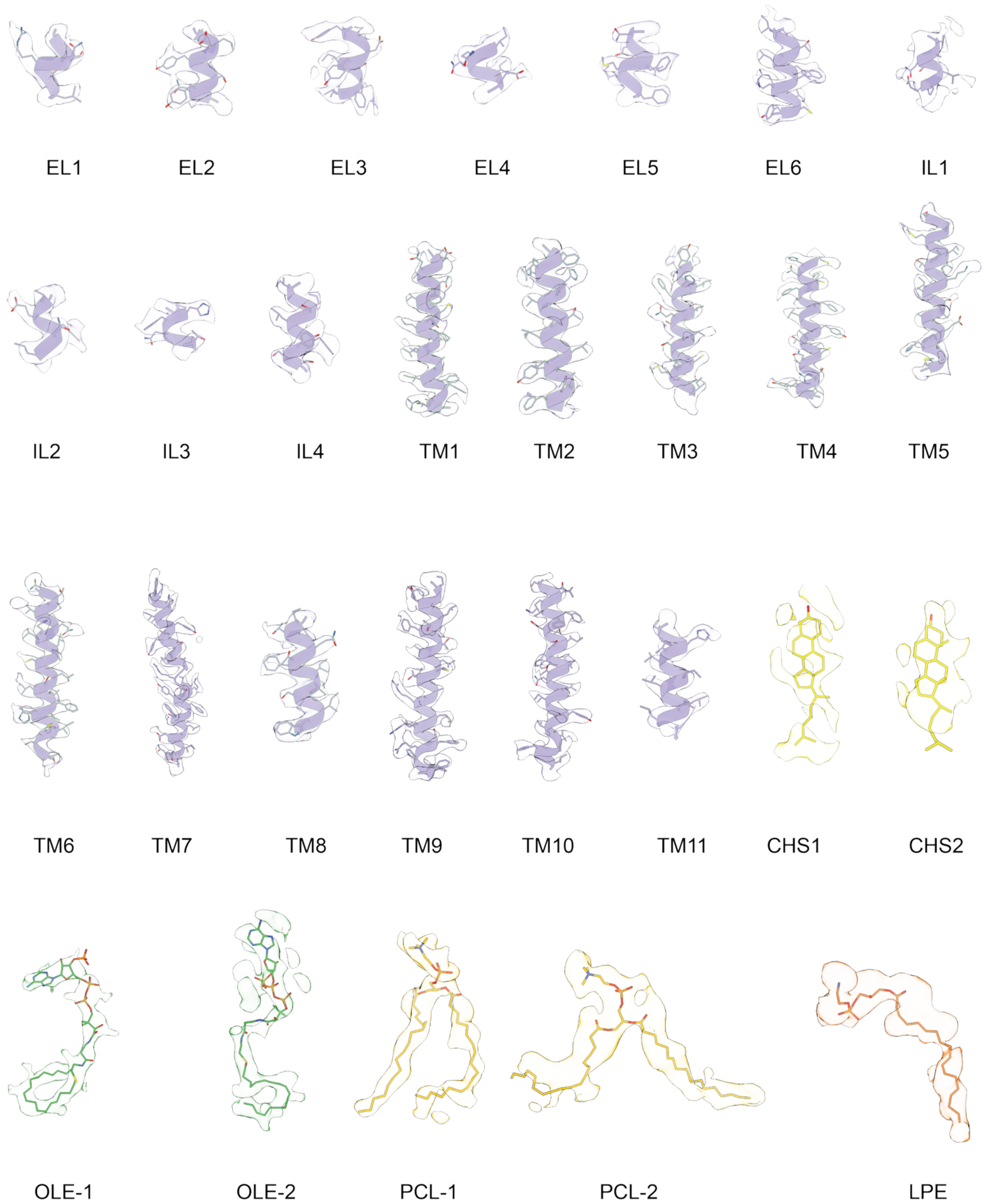


**Extended Data Fig. 9: EM density of different parts of human MBOAT2 in State D.**

### Extended Data Table 1 | Cryo-EM data collection, refinement and validation statistics

| | MBOAT2 apo (EMDB: pending) (PDB: pending) | MBOAT2 H373A monomer (EMDB: pending) (PDB: pending) | MBOAT2 oleoyl-CoA-added dimer (EMDB: pending) (PDB: pending) | MBOAT2 LPE-added dimer (EMDB: pending) (PDB: pending) |
|---|---|---|---|---|
| **Data collection and processing** | | | | |
| Magnification | 130,000 | 130,000 | 130,000 | 130,000 |
| Voltage (kV) | 300 | 300 | 300 | 300 |
| Electron exposure (e-/Å²) | 40 | 40 | 40 | 40 |
| Defocus range (μm) | −0.8 to 1.8 | −0.8 to 1.8 | −0.8 to 1.8 | −0.8 to 1.8 |
| Pixel size (Å) | 0.940 | 0.940 | 0.940 | 0.940 |
| Symmetry imposed | C2 | C1 | C2 | C2 |
| Initial particle images (no.) | 7,796 | 8,453 | 7,796 | 7,421 |
| Final particle images (no.) | 7,796 | 8,453 | 7,796 | 7,421 |
| Map resolution (Å) | 3.14 | 3.54 | 3.35 | 3.48 |
| FSC threshold | 0.143 | 0.143 | 0.143 | 0.143 |
| **Refinement** | | | | |
| Initial model used (PDB code) | AlphaFold | MBOAT2 apo model | MBOAT2 apo model | MBOAT2 apo model |
| Resolution (Å) | 3.14 | 3.54 | 3.35 | 3.48 |
| **Model composition** | | | | |
| Non-hydrogen atoms | 8,067 | 4,047 | 8,027 | 7,545 |
| Protein residues | 904 | 452 | 904 | 904 |
| Ligands | 20 | 9 | 16 | 8 |
| **B factors (Å²)** | | | | |
| Protein | 60.98 | 57.10 | 80.72 | 71.26 |
| Ligand | 78.83 | 65.69 | 66.71 | 100.64 |
| **R.m.s. deviations** | | | | |
| Bond lengths (Å) | 0.002 | 0.003 | 0.004 | 0.003 |
| Bond angles (°) | 0.647 | 0.640 | 0.930 | 0.790 |
| **Validation** | | | | |
| MolProbity score | 1.97 | 2.12 | 1.95 | 2.51 |
| Clashscore | 7.56 | 7.40 | 5.92 | 14.06 |
| Poor rotamers (%) | 2.13 | 3.77 | 2.26 | 4.02 |
| **Ramachandran plot** | | | | |
| Favored (%) | 95.65 | 95.98 | 94.87 | 94.20 |
| Allowed (%) | 4.35 | 4.02 | 5.13 | 5.80 |
| Disallowed (%) | 0.00 | 0.00 | 0.00 | 0.00 |
